# Adult facilitation becomes competition as juvenile soapberry bugs age

**DOI:** 10.1101/186536

**Authors:** Meredith L Cenzer

## Abstract

Whether intraspecific interactions are facilitative or competitive may change across individual ontogeny. In plant-feeding insects, the direction of this interaction is likely to be mediated by host plant defenses. Here I conducted two experiments looking at the direct effect of a physical seed defense and the role of intraspecific facilitation in reducing the effects of that defense for juveniles. I first demonstrate that juveniles of the red-shouldered soapberry bug (*Jadera haematoloma*) are severely inhibited by the tough seed coat of their host plant, leading to high mortality early in development. Adults, in contrast, can create holes through which other individuals could potentially feed. I then manipulated whether or not seeds experienced adult feeding on two host plant species: a well-defended native host, balloon vine (*Cardiospermum corindum*) and a poorly defended introduced golden rain tree species (*Koelreuteria elegans*). I measured the effect of prior adult feeding on survival, development time, and final body size of soapberry bug juveniles. Survival in the first week of development was dramatically improved by prior adult feeding on both hosts. However, the benefits of prior adult feeding ceased after the first week of development and shifted to having a negative effect on performance. These results indicate that adults breaking through the seedcoat initially facilitate juveniles, but that this facilitation becomes competition as juveniles age.

## INTRODUCTION

Plant defenses are important determinants of herbivore fitness that mediate indirect interactions between herbivores. In plant-feeding insects, host plant defenses are a common moderator of both competition (Karban and Baldwin 1997) and facilitation (Soler et al. 2012; Huang et al. 2013; Glas et al. 2014). Competition is the most common net outcome of multiple individual herbivores sharing a plant (Kaplan and Denno 2007), primarily via the induction of plant defenses (Karban and Baldwin 1997). Facilitation within or between herbivorous species has been shown via the suppression or modification of plant defenses (Soler et al. 2012; Huang et al. 2013; Glas et al. 2014).

The relative importance of plant defenses and resource limitation is likely to vary across individual ontogeny. As individuals of most species age, both their ability to access well-defended resources alone and the quantity of resources they require are likely to increase. Therefore, facilitative interactions may become competitive as individuals age. This pattern is commonly observed in plants (e.g., nurse plants), where it has important implications for population and community structure (Callaway and Walker 1997). In animals, there are many observations that support some part of this process, but cases of facilitation becoming competition across ontogeny have rarely been directly observed. For example, gregarious nymphalid caterpillars have higher weight gain when feeding in large groups in the field, but are also observed to move to mature leaves as they age due to defoliation of their host plants (Denno and Benrey 1997). Feeding on mature leaves has been shown to decrease growth rate and biomass conversion efficiency in other caterpillar species (Stamp and Bowers 1990), suggesting increased weight gain from group feeding early in life likely comes with a cost to growth rate later in development.

This has important ecological implications for how host use and behavioral interactions (e.g., aggregation/aggression) should change across development. For example, a shift from facilitation to competition has been suggested to explain why many group-feeding larval aggregations, which are regularly shown to have improved performance over solitary larvae, break up as juveniles age (Denno and Benrey 1997; Fordyce 2003; Inouye and Johnson 2005). Shifts in interaction direction also have important logistical implications about when and how we choose to study plant-herbivore interactions.

In this study, I evaluate the role of a host plant defense, the seedcoat, in juvenile performance across ontogeny. I then look at how prior adult feeding, which creates physical holes in the seedcoat, influences juvenile performance both early and late in development. I evaluate these effects on two host plants, one native and one introduced.

### Study system

The red-shouldered soapberry bug *Jadera haematoloma* (Hemiptera: Rhopalidae) is a seed-feeding specialist native to the southern peninsula and Keys of Florida, where it has evolved with a native balloon vine (*Cardiospermum corindum). Cardiospermum corindum* has seeds that are defended by an inflated seedpod, a hard seedcoat, and chemical toxins, especially saponins (Umadevi and Daniel 1991). Juveniles can never reach seeds inside of a closed seedpod, which serves primarily as a barrier to adult feeding (Cenzer 2017). Juveniles reared on dehisced seeds of this host have very high mortality during the first week of development (Cenzer 2016a), suggesting they are unable to circumvent the seed’s defenses at this stage. In contrast, adults and older juveniles have relatively low mortality on this host. After feeding, adults leave visible holes behind that may serve as access points for young individuals. However, these access points may come at the cost of losing nutrients to individuals who fed earlier, creating the possibility of resource limitation as juveniles grow and their resource demand increases. Soapberry bugs often feed together in mixed age groups, creating ample opportunities for both positive and negative interactions between age classes.

In the 1950s, a species of golden rain tree (*Koelreuteria elegans*) was widely introduced to the peninsula of Florida and colonized by soapberry bugs. Unlike the native *C. corindum, K. elegans* has relatively soft seeds that don’t present a strong physical barrier to feeding. Juvenile survival on *K. elegans* is quite high even when juveniles are reared alone, reducing the potential benefits of group feeding (Cenzer 2016b). In the 1980s and 1990s, local adaptation was documented between soapberry bug populations between the introduced and the native hosts in Florida in juvenile survival, development time, and feeding morphology (Carroll and Boyd 1992; Carroll et al. 1997, 1998) indicating that differences between these two host plants influence juvenile performance.

In this study, I address the questions: How does breaking through the seedcoat, both manually and via prior adult feeding, influence juvenile performance? How do these effects differ early and late in development? How does the effect of the seedcoat differ between the native and introduced host plants? I tested the following four hypotheses: H1) Manually cracking the physical barrier of the seedcoat has a positive effect on fitness. H2) Prior adult feeding facilitates juvenile performance by creating physical entry points in the seedcoat, increasing fitness. H3) The positive effects of prior adult feeding will be most pronounced early in development, when juvenile survival is lowest, and weaken later in development. H4) The benefits of penetrating the seedcoat barrier will be stronger on the more well-defended native host plant.

## METHODS

### Collection

I collected adult *J. haematoloma* for Experiment 1 in April 2014 from 7 locations in Florida: Gainesville [*K. elegans*], Leesburg [*K. elegans*], Lake Wales [*K. elegans*], Ft. Myers [*K. elegans*], Homestead [*K. elegans*], Homestead [*C. corindum*], Key Largo [*C. corindum*], and Plantation Key [*C. corindum].* For Experiment 2, I collected in March-April 2015 from four of these locations in Florida (Leesburg [*K. elegans*], Lake Wales [*K. elegans*], Key Largo [*C. corindum*], Plantation Key [*C. corindum]*) and one location in California (Davis [*K. paniculata*])(Appendix A). I collected host plant seeds from each Florida site in December 2013, April 2014, and April 2015 and stored them at 4°C until they were used for rearing. Seeds were only collected from *K. paniculata* in April 2015 (Appendix A). Seeds with visible indications of previous feeding were discarded. I tested all seeds for viability by placing them in water and discarding seeds that floated. I collected from 5-10 individual trees at each *K. elegans* site, 3-15 individual vines at each *C. corindum* site, and from 6 trees at the *K. paniculata* site.

### Experiment 1: Manually cracking the seedcoat

The first experiment tested the effects of the seed coat on juvenile survival. All rearing was carried out in controlled environmental chambers (Sanyo Versatile Environmental Test Chambers) at 28°C during the day and 27.5°C at night, 50% relative humidity with a 14:10 light:dark cycle, following spring climate conditions in the field and those used in earlier work (Carroll et al. 1998; Cenzer 2016b). Adults collected from the field in April 2014 were housed as mating pairs in vented Petri dishes lined with filter paper and given water in a microcentrifuge tube stoppered with cotton (“water pick”) and 3 seeds of their field host plant. For this experiment, I used the second laboratory generation of juveniles descended from the April 2014 field collection. The first laboratory generation was used for experiments described in Cenzer 2016. Eggs were collected daily until hatching. Nymphs were removed within 12 hours of hatching to reduce egg cannibalism and housed individually in mesh-lidded cups lined with filter paper with a water pick and a seed of their assigned rearing host treatment.

Juveniles were distributed in a split-brood cross-rearing design, such that full siblings from all families were represented in all treatments. Upon hatching, juveniles were randomly assigned to a rearing host (either *C. corindum* or *K. elegans*) and a seed treatment (intact or cracked seedcoat). I administered the cracked seed treatment by gently clamping seeds in pliers and tightening the pliers just until a crack formed in the seedcoat. Seven days after hatching, additional seeds (a total of 2 for *K. elegans* and 3 for *C. corindum*, for a total seed mass of ~150mg) were added to each juvenile’s container. Individual containers were rotated daily within mesh boxes, and boxes were rotated daily within the growth chamber. Water, paper and cotton were changed weekly. Nymph survival and whether or not they had reached adulthood was assessed daily. Upon reaching adulthood, bugs were allowed 1 day for the exoskeleton to harden and were then frozen at −20°C for morphological analyses.

### Experiment 1: Statistical analyses of the fitness effects of the seedcoat

All analyses were conducted in R version 3.3.3 (“Another Canoe”). The sets of models evaluated for each response variable are discussed in greater detail below. For each response variable, all models were compared using relative probability based on the Akaike Information Criterion (AIC, a metric that ranks the relative quality of a set of models based on fit and simplicity). All models with a >5% probability of being the best model out of the set were examined for each response variable. If effects were not significant and in the same direction in all examined models, models were directly compared using Chi-squared tests. The specific test statistics and effect sizes reported in the results section were taken from the model with the highest probability.

The performance cost of the seed coat (H1) was evaluated by including the seed treatment (intact or cracked) as a fixed factor in the analyses of survival, development time, and final adult body size. To evaluate whether the effects of the seed treatment on survival differed early and late in development, survival was subset into the first week of development (“early”) and after the first week (“late”). For total, early, and late survival, all possible models including the main effects of rearing host (*C. corindum* or *K. elegans*), ancestral host (*C. corindum* or *K. elegans*) seed treatment (cracked or intact), and all possible two-way interactions were considered. These models were also analyzed as generalized linear mixed models with the random factor of individual population nested within ancestral host and the random factor of family nested within individual population. Survival was modeled using a binomial distribution.

The response variables development time and body size had reduced sample sizes in some treatments due to strong treatment effects on survival. Therefore, only a subset of models were considered for these variables. For development time, all models with the main effects of seed treatment, rearing host, ancestral host, and sex and the rearing host * seed treatment interaction were compared. For body size, which is highly dimorphic between sexes, models with pairwise interactions with sex were also considered. For development time, I used a Gamma error distribution. Gamma distributions are positive definite with skew to the right; they are flexible distributions commonly used in modeling waiting times. I modeled body size with a Gaussian error distribution and tested residuals for normality using Shapiro-Wilk normality tests.

### Experiment 2: Prior adult feeding treatment

The second experiment tested whether prior feeding by adults could mimic the effects of cracking the seedcoat barrier. The two seed treatments were intact seeds and seeds that had experienced prior feeding by adults.

To produce the prior feeding treatment, I used adult soapberry bugs of mixed sex collected in Davis, CA from *K. paniculata.* To ensure that this population and host were not unusual, this host was included as both an ancestral and rearing host (see Appendix A). The results of these treatments did not differ qualitatively from those on the other two hosts, and there was no evidence of local adaptation to *K. paniculata*, indicating that adults from this host are unlikely to feed differently than adults from any other host would. Adults were held for 24 hours without food to ensure hunger. Seeds of all three species were soaked for 18 hours to encourage adult feeding. Vented Petri dishes were set up with filter paper, a water pick, and two seeds of different species. Seeds were secured in place in each dish using modeling clay. Five adults of mixed sex were introduced into dishes assigned to the prior feeding treatment; dishes assigned to the intact treatment had no adults. Petri dishes were returned to the incubator and remained for an average of 55 hours before being used for rearing. This method resulted in 94.7-98.9% of seeds assigned to the prior feeding treatment receiving visible feeding damage. If a seed was assigned to the prior feeding treatment and did not show visible signs of feeding, it was not used in the experiment. This exclusion may have created a bias such that very poor quality seeds might not have been used in the prior feeding treatment due to filtering by adult choice. Comparison with field measures of hole number on *C. corindum* indicate that the amount of damage induced by this treatment represents the lower end of the range of damage in the field (Fig. 1). The availability of seeds per bug was less than one (unpublished data) on *C. corindum*, so seed availability in this experiment was generous. Because *K. elegans* produces a single massive crop of seeds in the late fall that is completely depleted by April (Carroll et al. 1998), the per-seed damage on this host varies continuously over the year from no damage on any seeds to complete removal of all resources and is not well represented by a point sample.

**Figure 1.**
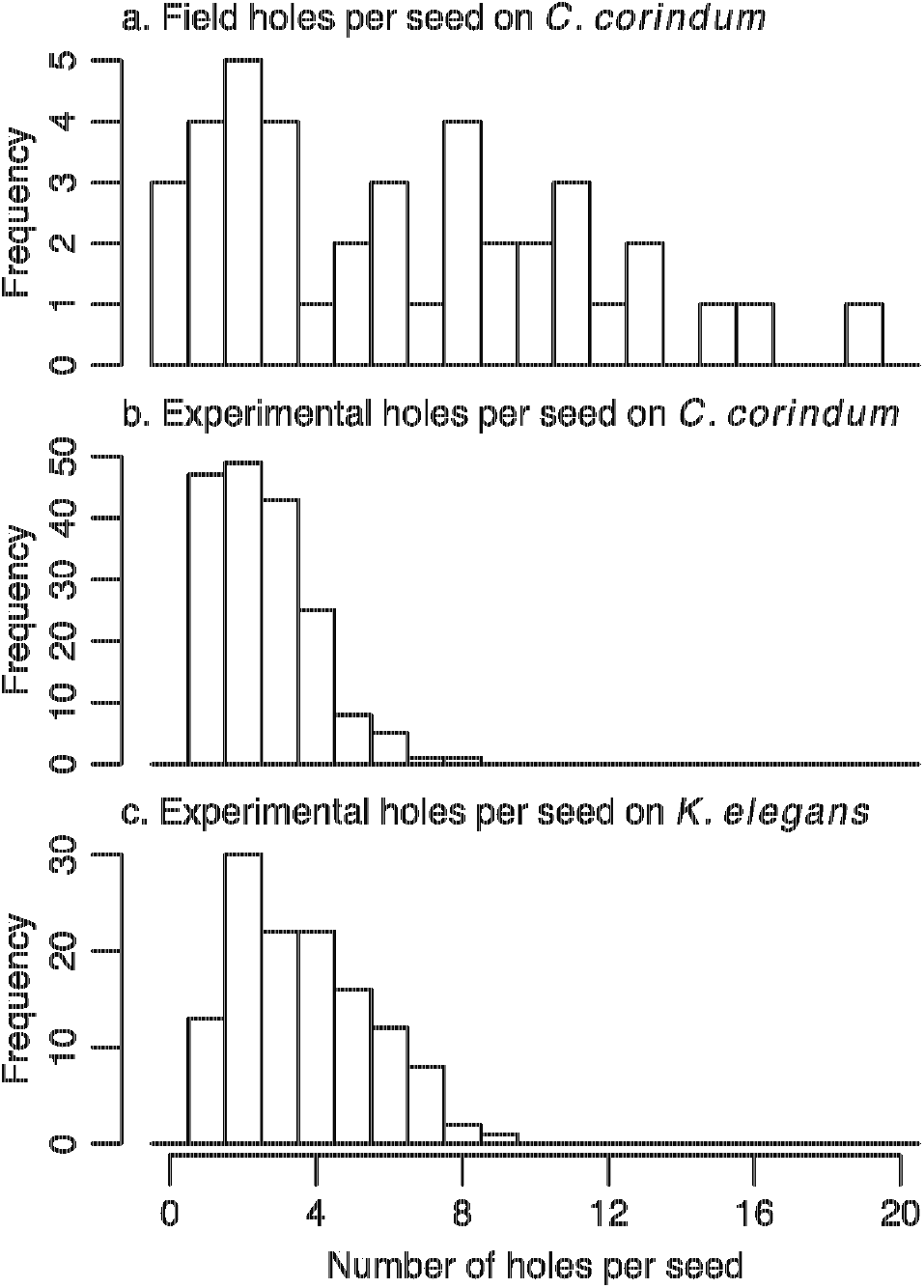
The number of holes per seed measured in the field on *C. corindum* (a) and experimentally produced in the prior feeding treatment on both *C. corindum* (b) and *K. elegans* (c) in Experiment 2. Seeds with no feeding damage are included in field measures but are not included in experimental measures.

For this experiment, juveniles were used from the first laboratory generation descended from the 2015 field collection. Juveniles were reared following the same protocol as described above in Experiment 1, in two rearing host treatments (*C. corindum* and *K. elegans*) and two seed treatments (intact and prior feeding). The number of holes in each seed was recorded before each juvenile began feeding.

### Experiment 2: Statistical analyses of the fitness effects of prior adult feeding

Model selection and reporting follow the same procedures as described above for Experiment 1.

Total and early survival, development time, and body size were all analyzed using the same sets of models considered above in Experiment 1. Early survival was substantially lower in some treatments in this experiment, and late survival was therefore analyzed using a subset of models for total survival. Models containing all possible combinations of the main effects of seed treatment, rearing host, and ancestral host, as well as the rearing host * seed treatment interaction were compared.

In order to address the hypothesis that changes in survival, development time, and body size were the result of physically breaking through the seed coat, I also tested the effect of the number of holes drilled by adults on each of these response variables. Analyses of hole number were run only on data from the prior feeding treatment to distinguish the main treatment effect from the effect of hole number. This effect was evaluated using the same models that were considered for each response variable on the full dataset, but replacing seed treatment with hole number.

It should be noted that variation in hole number was produced by adult feeding decisions rather than by direct manipulation. Therefore, analyses of hole number may be testing the effect of some underlying variable that influenced the number of holes adults chose to drill in each seed (ie, seed toughness) rather than the direct effect of hole number itself.

## RESULTS

### H1) Cracking the seedcoat increased juvenile survival

In Experiment 1, the cracked seed treatment produced a dramatic increase in survival probability on both *C. corindum* (0.079 vs 0.94; z-value=7.70, p<0.001) and *K. elegans* (0.83 vs 0.98; z-value=2.39, p=0.017).

Nymph mortality was heavily skewed towards very young individuals, such that 94% of nymph mortality occurred in the first 7 days after hatching (Fig. S1). During this early period, results paralleled those for total survival. The cracked treatment increased early survival on both *C. corindum* and *K. elegans* (z-values=5.91, 2.12; p<0.001, p=0.034) (Fig. 2a). The cracked treatment continued to have a positive effect on survival late in development (z-value=3.19, p=0.001) (Fig. 2c) that did not differ detectably between host plants.

**Figure 2.**
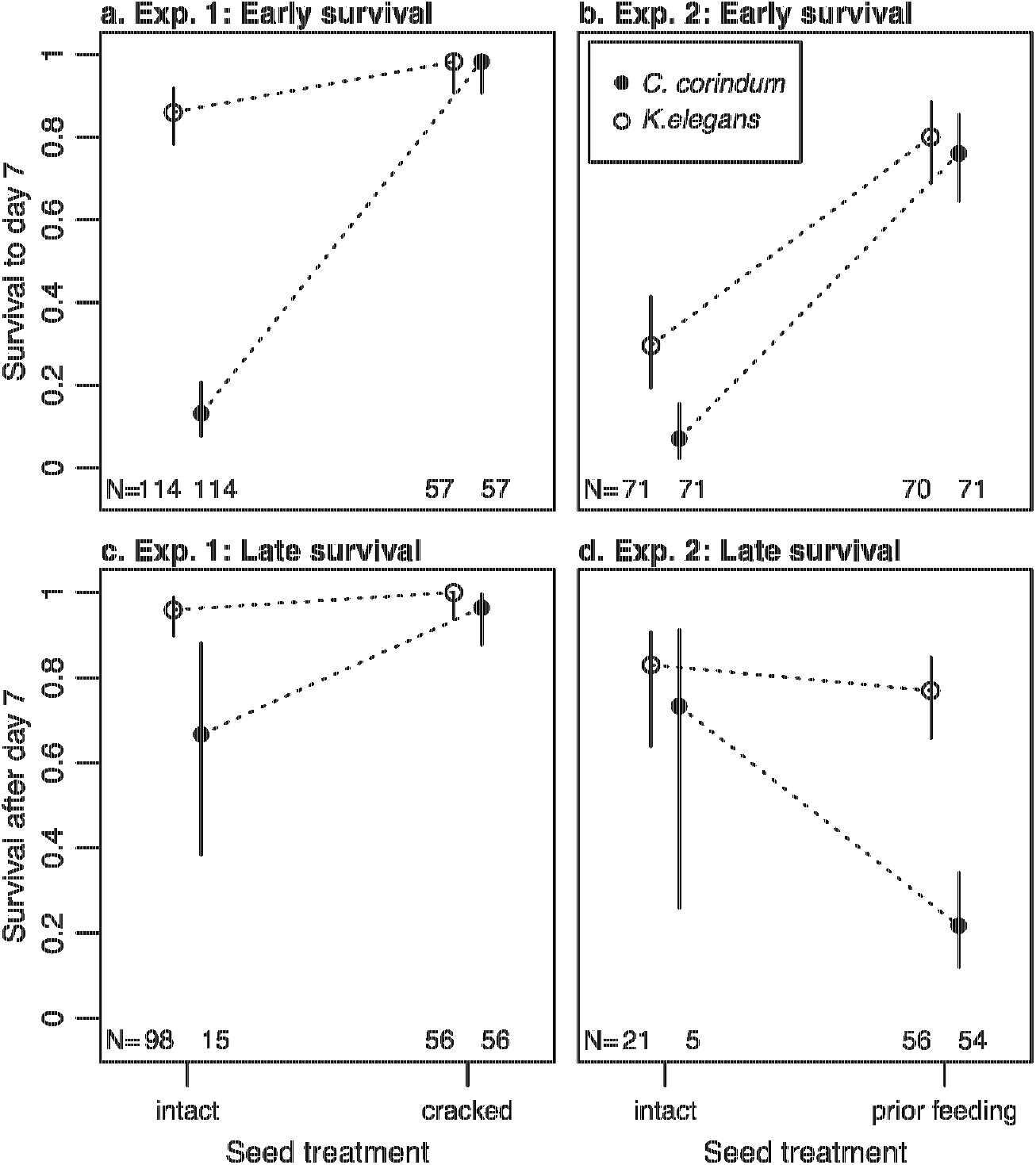
Soapberry bug juvenile survival before (a) and after (b) day 7 in Experiment 1 (squares) and Experiment 2 (circles). Left-hand panels are bugs reared on host *C. corindum* and right-hand panels are bugs reared on *K. elegans.* The seed manipulation in Experiment 1 is cracked; in Experiment 2 it is the natural analog, prior adult feeding. Points are means and error bars are 95% binomial confidence intervals computed using the Pearson-Klopper method.

**Figure 3.**
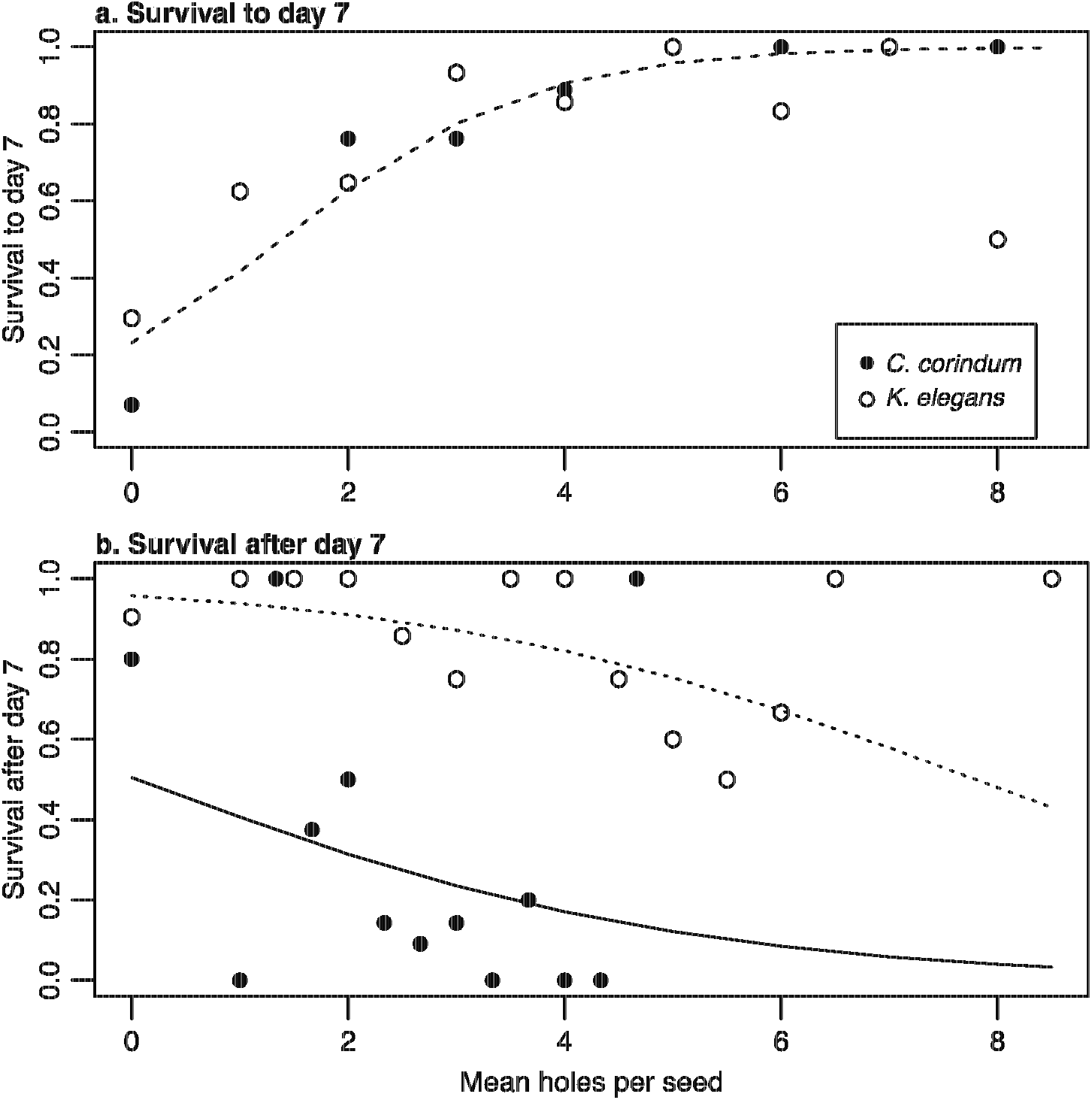
Survival prior to day 7 (a) and after day 7 (b) as a function of the number of holes drilled by adults. All intact seeds have 0 holes. Rearing hosts are *C. corindum* (filled circles) and *K. elegans* (open circles). In (a), the dashed line shows model predictions for the relationship between hole number and survival up to day 7; this relationship did not differ between rearing hosts. In (b), the solid line shows model predictions for the relationship between hole number and survival after day 7 on rearing host *C. corindum* and the dotted line shows model predictions on rearing host *K. elegans.*

On *C. corindum*, juveniles in the cracked seedcoat treatment developed more quickly than those on the intact treatment (t-value=-2.65, p=0.01, df=199); cracking the seedcoat was neutral on *K. elegans* (t-value=-1.28, p=0.2)(Fig. 4a). Juveniles in the cracked and intact seed treatments did not differ in their final adult body size (Fig. 5a).

**Figure 4.**
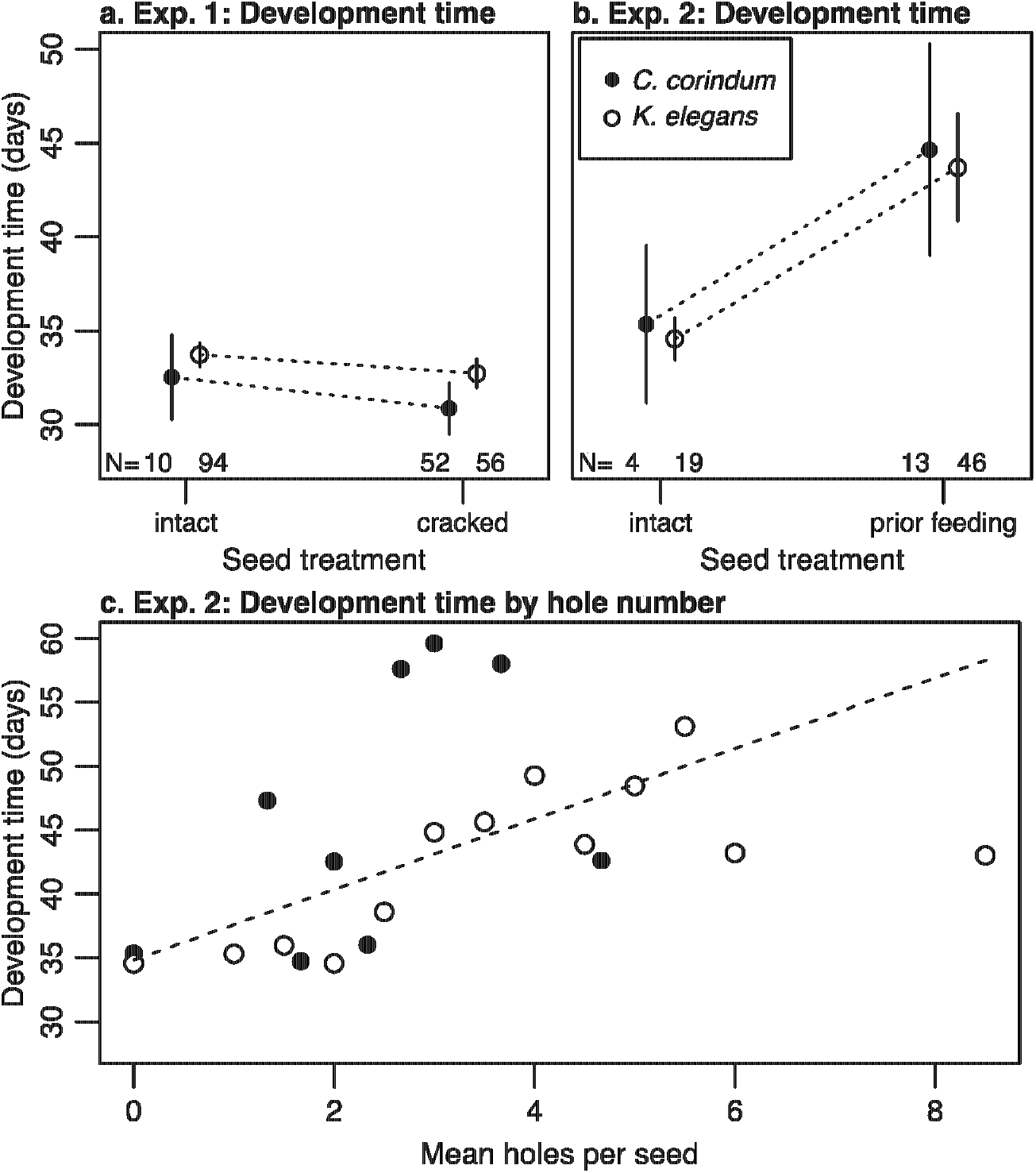
Development time from hatching to adulthood by treatment. All development times are corrected for ancestral host and sex. a. Mean development times for bugs in Experiment 1 on intact and cracked seeds for bugs reared on *C. corindum* (circles) and *K. elegans* (squares). b. Mean development times for bugs in Experiment 2 on intact and prior feeding treatment seeds for bugs reared on *C. corindum* (circles) and *K. elegans* (squares). c. Development time as a function of the number of holes in each seed for bugs reared on *C. corindum* (filled circles) and *K. elegans* (open circles). The dashed line represents model predictions from the top model, which did not differ between rearing hosts.

**Figure 5.**
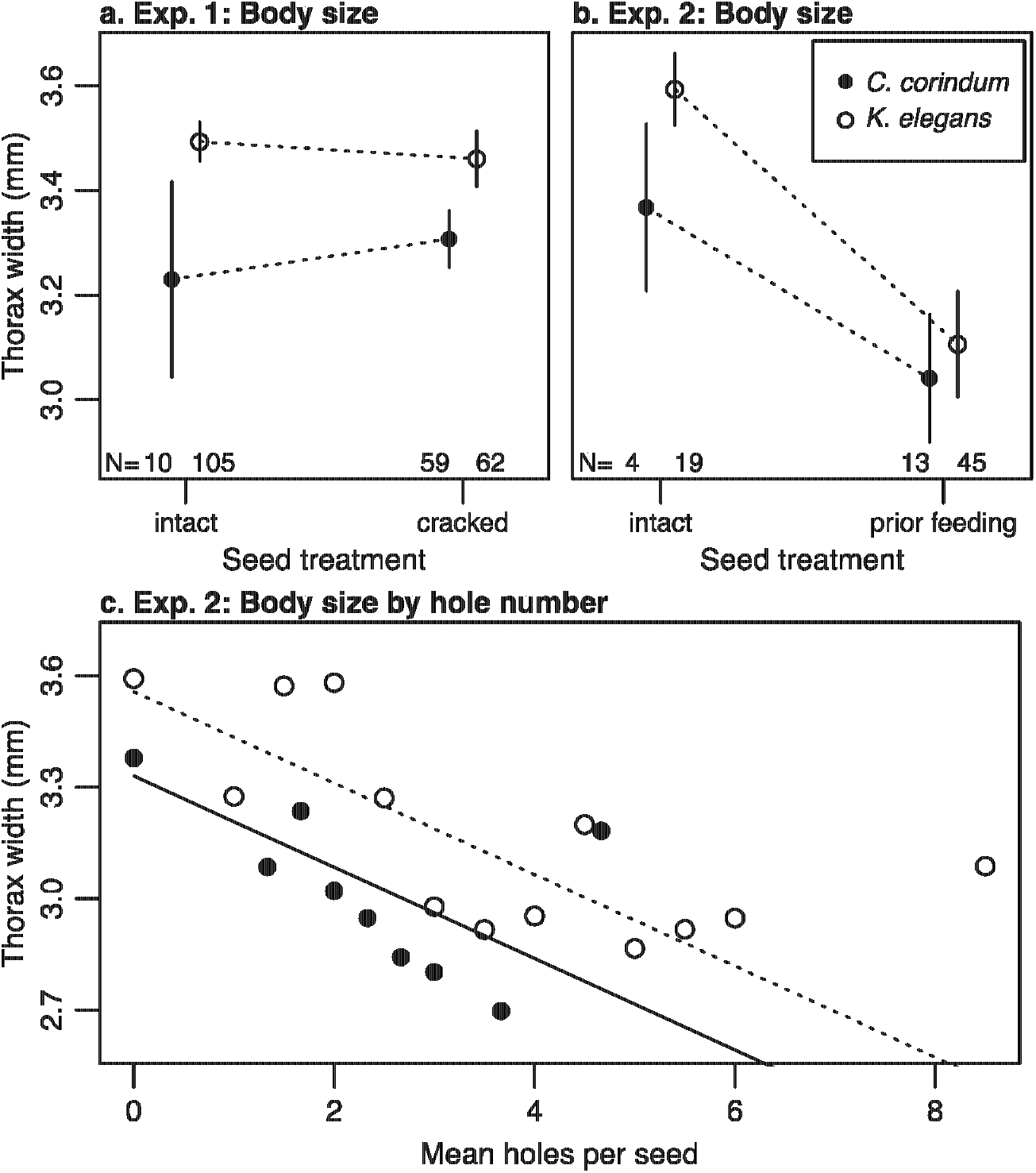
Adult thorax width by treatment. All measures are corrected for sex. a. Mean adult thorax width (mm) in Experiment 1 on rearing hosts *C. corindum* (circles) and *K. elegans* (squares) on intact and cracked seeds. Error bars represent 95% confidence intervals. b. Mean adult thorax width (mm) in Experiment 2 on rearing hosts *C. corindum* (circles) and *K. elegans* (squares) on intact and prior feeding treatment seeds. c. Thorax width as a function of the number of holes in each seed for bugs reared on *C. corindum* (circles, solid line) and *K. elegans* (squares, dashed line). Lines represent model predictions from the top model for each rearing host.

### H2) Early in development, prior adult feeding mimics the positive effects of cracking the seedcoat

Results for early survival in Experiment 2 parallel those in Experiment 1. Juveniles in the prior feeding treatment had a significantly higher probability of surviving to adulthood than juveniles on the intact seed treatment on both rearing hosts (z-value=5.18, p<0.001). This effect was produced by increased early survival in the prior feeding treatment on both *C. corindum* and *K. elegans* (z-values=6.94, 5.72; p<0.001)(Fig. 2b). Consistent with a shared mechanism between Experiments 1 and 2, there was a strong and positive effect of the number of holes drilled by adults in each seed on survival to day 7 in the prior feeding treatment (z-value=2.15, p=0.03) (Fig. 3a).

### H3) Late in development, prior adult feeding has strong negative effects on performance

In contrast, the effects of the prior feeding treatment later in development were no longer positive. Bugs reared in the prior feeding treatment suffered increased late mortality when reared on *C. corindum* (z-value=2.32, p=0.02) and gained no benefit when reared on *K. elegans* (z-value=0.43, p=0.6) (Fig. 2d). Increasing the number of holes within the prior feeding treatment had a strong negative effect on survival after day 7 independent of rearing host (z-value=-2.27, p=0.02) (Fig. 3b).

The prior feeding treatment also increased the amount of time it took to reach adulthood (t-value=-4.18, p<0.001, df=78) (Fig. 4b). Consistent with the treatment effect, each additional hole increased development time within the prior feeding treatment (t-value=3.60, p<0.001, df=56) (Fig. 4c).

Females and males in the prior feeding treatment were both significantly smaller than those in the intact treatment (t-values=-6.06, −3.27; p<0.01; df=76) (Fig. 5b). Within the prior feeding treatment, increasing the number of holes in a seed decreased adult body size (t-value=-3.12, p=0.003, df=55) (Fig. 5c).

### H4) Juveniles reared on the native host were more sensitive to benefits and costs of penetrating the seedcoat

The seedcoat had a stronger effect on early juvenile survival when reared on the native host than on the invasive host. In Experiment 1, there was a clear difference in survival between the two rearing host plants in the intact seed treatment (z-value=9.43, p<0.001) that vanished when seeds were cracked (z-value=0.097, p=0.33).

Similar to results in Experiment 1, the seed treatment in Experiment 2 eliminated differences in early survival between hosts. Without prior feeding, individuals reared on *K. elegans* had higher early survival than those on *C. corindum* (z-value=3.22; p=0.001), but this difference vanished in the prior feeding treatment (z-value=-0.56; p=0.58).

Juveniles on the native host also benefited more in terms of development time. On *C. corindum*, juveniles in the cracked seedcoat treatment developed more quickly than those on the intact treatment (t-value=-2.65, p=0.01, df=199); cracking the seedcoat had no effect on development time on *K. elegans* (t-value=-1.28, p=0.2)(Fig. 4a).

Juveniles reared on the native host also suffered increased costs of prior adult feeding compared to those on the introduced host. Bugs reared in the prior feeding treatment suffered increased mortality late in development when reared on *C. corindum* (z-value=2.32, p=0.02), but not on *K. elegans* (z-value=0.43, p=0.6) (Fig. 2d). Being reared on *K. elegans* had a positive overall effect on late survival (z-value=5.83, p<0.001) and a marginally positive effect on adult body size (t-value=1.93, p=0.059, df=55).

## DISCUSSION

I found strong support for my first hypothesis that the tough seedcoat of the host plant has a strong negative effect on juvenile fitness, primarily by decreasing survival. I also found support for my second hypothesis that prior adult feeding is a natural analog of cracking the seedcoat early in development, facilitating early juvenile survival by creating physical access points through the seed coat. Later in development, cracking the seedcoat continued to have positive or neutral effects on juvenile fitness. After the vulnerable first week of development, however, the effects of prior adult feeding not only weakened, as I predicted in my third hypothesis, but became strongly negative. I found that early facilitation trades off with the cost of increasing mortality late in development, prolonging development time, and decreasing adult body size, likely due to direct competition for resources. I also found that bugs reared on the native host plant, *C. corindum*, were more sensitive to the positive and negative effects of breaking through the seedcoat barrier than the introduced *K. elegans.*

I demonstrate here that the seedcoat is the proximate driver of early juvenile mortality. Early mortality for individuals raised on intact seeds is probably the result of newly hatched individuals being unable to penetrate the seed and subsequently starving to death. This mechanism was clearly paralleled by the effects of prior adult feeing, which had a positive net effect on juvenile survival on both rearing hosts. During this early developmental period, the number of holes drilled by adults was correlated with increased survival, consistent with the mechanism of increased survival being increased access to the endosperm within the tough seed. However, because hole number was determined by feeding adults and not manipulated directly, it is possible that both adult feeding frequency and juvenile survival were influenced by some unmeasured quality of the seed. Chemical defenses within the seed’s exterior, rather than or in addition to the physical barrier, may also contribute to early mortality.

After this critical early period in development, the effect of cracking the seedcoat on survival became weaker, although still positive. However, prior adult feeding ceased to parallel this positive effect, and instead had the opposite effect of decreasing survival late in development. The number of holes created by adults within the prior adult feeding treatment flipped to having a negative effect on juvenile survival late in development, likely because more intense adult feeding removed more nutrients from the seeds, resulting in stronger competition and eventual starvation for juveniles with seeds that experienced heavy feeding. This suggests that, in nature, there is a benefit for juveniles in having adults present because they allow easier access to seeds; however, as juveniles age, this benefit may be outweighed by the costs of competition.

Cracking the seedcoat manually had little effect on development time or final body size. However, the prior feeding seed treatment came at the cost of increased development times and smaller body sizes. Both of these costs were correlated with the increasing number of holes in each seed, again suggesting that increasing facilitation early in life came at a direct cost to increasing competition later in development. The difference in development time and body size between seed treatments was likely the result of partial starvation in the prior feeding treatment due to the removal of nutrients by prior feeding adults, such that individuals were not able to achieve the optimal size for eclosion before molting. This hypothesis is supported by the fact that the number of holes, and therefore prior adult feeding events, on each seed was correlated with increased development time and decreased body size. Consistent with nutrient deprivation, individuals who did not reach adulthood - and therefore had no measured development time or adult size - often persisted as nymphs well beyond the average time to adulthood and died, on average, at a much later age (Fig. S2). The cost of prior adult feeding could potentially be reduced in nature in some cases where individuals would be able to actively search for additional seeds, although field estimates of damage and seed availability are not promising on the native host (Fig. 1a) or late season on the introduced host (Carroll et al. 1998). Juveniles in nature also face continuous competition from both adults and other juveniles.

I also found that the costs of the seedcoat, and subsequently the benefits of breaking through the seedcoat, were stronger on the native *C. corindum* than on the introduced *K. elegans.* Although survival was generally higher on *K. elegans*, cracking the seedcoat and prior feeding both eliminated this difference early in development. Cracking the seedcoat also had a positive effect on development time on *C. corindum* that was absent on *K. elegans.* However, the negative effects of prior feeding on survival late in development were also more pronounced on *C. corindum* than on *K. elegans.* This difference may be due in part to the difference in seed size on these two hosts - *C. corindum* has smaller seeds that may be more rapidly depleted of resources once feeding has begun. This may effectively create a narrower window in which juveniles can access seeds on *C. corindum* after adults have begun feeding but before all resources have been removed from the seed.

The combined effects of adult feeding on juvenile survival may regulate population sizes. At high adult densities (relative to seed availability), seeds should be quickly depleted of resources, limiting nutrient availability for juveniles, who can rarely gain the benefit of early access to undamaged seeds. At very low adult densities, in contrast, most seeds will not have feeding damage and will therefore be unavailable as a resource for young juveniles. For example, if an adult female arrived on an unoccupied host plant, reproduced, and died without creating appreciable damage to seeds on a plant, her offspring would likely suffer high mortality from being unable to feed on local seeds. This has different implications for each host plant in nature. The native *C. corindum* has smaller populations and more rapid patch turnover due to asynchronous fruit production; patches are regularly recolonized and may have low initial densities. The high mortality induced by the seedcoat of this host could therefore be further exacerbated in nature by the fact that juveniles will often have to cope with seed defenses in the absence of much adult facilitation. There are, however, other species of seed predators on the native host (eg, the silver-banded hairstreak *Chlorostrymon simaethis* [Drury 1773]) that damage the seedcoat, and could plausibly facilitate juvenile survival in the absence of adult soapberry bugs. In contrast, the invasive host, *K. elegans*, supports very large populations of soapberry bugs in the field that persist throughout the season and overwinter on or near the host, increasing the potential for both intraspecific facilitation and competition. Individuals of *K. elegans* produce tens of thousands of seeds each year (Carroll et al. 1998), which may buffer intraspecific competition until late in the season, when competition is likely to become intense as resources are depleted prior to summer diapause.

Together, these two experiments indicate that adults facilitate juvenile soapberry bugs early in development via the breakdown of a physical plant defense, the seedcoat. The benefits of facilitation are substantial but short-lived: once juveniles grow out of the vulnerable first instar, there is no longer a measurable benefit to using seeds with prior feeding. I also found that juveniles that develop on the native host plant are more sensitive to the effects of prior feeding, suggesting bugs on this host are both more reliant on and have a narrower window of opportunity for facilitation in nature. These results suggest very young juveniles should seek out seeds with prior feeding, while older juveniles should preferentially use pristine seeds. This may create a conflict between developmental stages, as older juveniles make seeds accessible to younger juveniles, and should promote behavioral changes across ontogeny from aggregative to more solitary. This pattern of behavior is often observed in other insect herbivores, and highlights the importance of ontogeny in the study of plant-herbivore interactions.

## ACKNOWLEDGEMENTS

I would like to thank LH Yang, SP Carroll, JA Rosenheim, and J Mutz for comments on earlier versions of this manuscript. I would also like to thank the Florida Division of Recreation and Parks for permission to collect *Jadera haematoloma* and *Cardiospermum corindum* seeds within Florida state parks. I am particularly indebted to Aarti Sharma for her essential role during the methods development of the adult facilitation experiment and assistance with conducting both experiments. Funding was provided by the University of California, Davis Department of Entomology, the Center for Population Biology, and a Henry A. Jastro Research Fellowship

